# Infants’ attention to eyes is an independent, heritable trait predicting later verbal competence

**DOI:** 10.1101/2021.10.23.465424

**Authors:** Charlotte Viktorsson, Ana Maria Portugal, Danyang Li, Maja Rudling, Monica Siqueiros Sanchez, Kristiina Tammimies, Mark J. Taylor, Angelica Ronald, Terje Falck-Ytter

## Abstract

From birth, infants orient preferentially to faces, and when looking at the face, they attend primarily to eyes and mouth – two areas that convey different types of information. Here, in a sample of 535 5-month-old infant twins, we assessed eye (relative to mouth) preference in early infancy. Eye preference was independent from all other concurrent traits measured, and had a moderate-to-high contribution from genetic influences (A = .57; 95% CI: .45, .66). Preference for eyes over mouth at 5 months predicted higher parent ratings of verbal competence in toddlerhood, but did not predict autistic traits. These results suggest that variation in eye looking reflects a type of biological niche picking emerging before infants can select their environments by other means (crawling or walking).

## Introduction

Looking behavior is important in infants’ interactions with their surrounding world, as it is the earliest capacity to act on the environment by discriminating and selecting inputs for learning (Amso & Scerif, 2015; Conejero & Rueda, 2017; Hendry, Johnson, & Holmboe, 2019). Earlier research shows that children at a very young age preferentially attend to social stimuli such as faces (Farroni, Menon, & Johnson, 2006), and that upright faces are scanned more extensively than both inverted and phase-scrambled faces (Gliga, Elsabbagh, Andravizou, & Johnson, 2009). Infants’ preferential looking at faces in complex displays increase considerably over the first year of life, while physical salience has a decreasing influence on where infants look (Frank, Amso, & Johnson, 2014; Frank, Vul, & Johnson, 2009; Kwon, Setoodehnia, Baek, Luck, & Oakes, 2016).

Different areas of the face convey partly different types of information. While the eyes transmit a range of socio-communicative and emotional information (Calder et al., 2002), the mouth is more strongly associated with visual speech information (Yehia, Rubin, & Vatikiotis-Bateson, 1998). Infants gradually change their preferential attention from the eyes to the mouth during the first year of life (de Boisferon, Tift, Minar, & Lewkowicz, 2017; Wagner, Luyster, Yim, Tager-Flusberg, & Nelson, 2013). Further, infants’ attention to the mouth increases when stimuli involve speech (Frank, Vul, & Saxe, 2012; Lewkowicz & Hansen-Tift, 2012; Tenenbaum, Shah, Sobel, Malle, & Morgan, 2013) and earlier studies indicate that a preference for the mouth from 6 months of age is associated with larger vocabulary (Tenenbaum et al., 2015; Tsang, Atagi, & Johnson, 2018; Young, Merin, Rogers, & Ozonoff, 2009).

A recent study has suggested substantial heritability of eye and mouth preference in toddlers aged 18-24 months (Constantino et al., 2017). However, to the best of our knowledge, no study has yet investigated the genetic and environmental contribution to eyes versus mouth preference in the first year of (postnatal) life, when visual input is critical for shaping brain development, and before infants can actively select their environment by other means (crawling or walking). Therefore, the primary aim of this study was to establish the relative role of genetic and environmental influences on eye preference (relative to mouth) in early infancy, using eye tracking. Because infants tend to look at either the eyes or the mouth when viewing faces, we operationalized viewing preference as a single measure of total looking time at the eyes relative to the total looking time at both the eyes and the mouth. Given the high monozygotic twin concordances found in the previous study of eye and mouth preference in toddlers (Constantino et al., 2017), we expected to find a moderate-to-high genetic contribution in young infants. This would provide evidence for active niche picking very early in life, whereby selective visual attention on aspects of the others’ faces is a heritable trait, which may shape subsequent learning and development (Kennedy et al., 2017).

Given the link between attention to the mouth and language development, we expected a positive association between a preference for looking at the mouth and follow-up measures of early language skills. Further, there is substantial, although inconsistent, evidence of atypical face scanning in children and adults with ASD, and we therefore tested whether there is an association between eyes relative to mouth preference and later socio-communicative difficulties. We did not, however, have a directional hypothesis in this case, due to conflicting results in the literature (Chita-Tegmark, 2016; Falck-Ytter & von Hofsten, 2011; Jones & Klin, 2013). ASD can be seen as the extreme end of a continuum spanning the whole population (Robinson et al., 2011; Robinson et al., 2016; Ronald et al., 2006), therefore, studies of autistic traits are relevant also for our understanding of autism as a clinical condition, and vice versa. All abovementioned hypotheses were pre-registered in OSF (https://osf.io/s8y74/). Informed consent was obtained from the parents of all the twins who participated. The study was approved by the regional ethics board in Stockholm and was conducted in accordance with the Declaration of Helsinki.

## Methods

### Participants

The Babytwins Study Sweden (BATSS) consisted of 622 same-sex twins (311 pairs) that were recruited from the national population registry (only the greater Stockholm area was selected). In total, 29% of the invited families participated in BATSS. Data collection was performed at the Centre of Neurodevelopmental Disorders at Karolinska Institutet (KIND) in Stockholm, Sweden. Sample demographics are fully reported elsewhere (Falck-Ytter et al., 2021).

General exclusion criteria for the study were opposite-sex twin pairs, diagnosis of epilepsy, known presence of genetic syndrome related to ASD, uncorrected vision or hearing impairment, very premature birth (prior to week 34), presence of developmental or medical condition likely to affect brain development (e.g., Cerebral Palsy, hydrocephalus), and infants where none of the biological parents were involved in the infant’s care. Among the recruited and tested infants, 3 twins were excluded from analysis because they subsequently were found not to fulfil the general criteria (above) due to seizures at the time of birth (n = 2 twins) and spina bifida (n = 1 twin). In addition, for this analysis we excluded infants due to twin-to-twin transfusion syndrome (n = 12 pairs), birthweight below 1.5 kg (n = 1 twin), and non-Swedish-speaking parents (n = 1 pair). Further, some infants did not provide any data due to technical reasons (n = 3 pairs), lack of time (n = 3 pairs + 1 twin), lack of room (n = 1 pair), infant being too tired or too fuzzy (n = 4 twins), and infant not having enough valid data for the task (38 twins, see section *Eye tracking* for details). The final sample consisted of 535 infants (see **Table 1** for descriptive statistics).

**Table 1.**
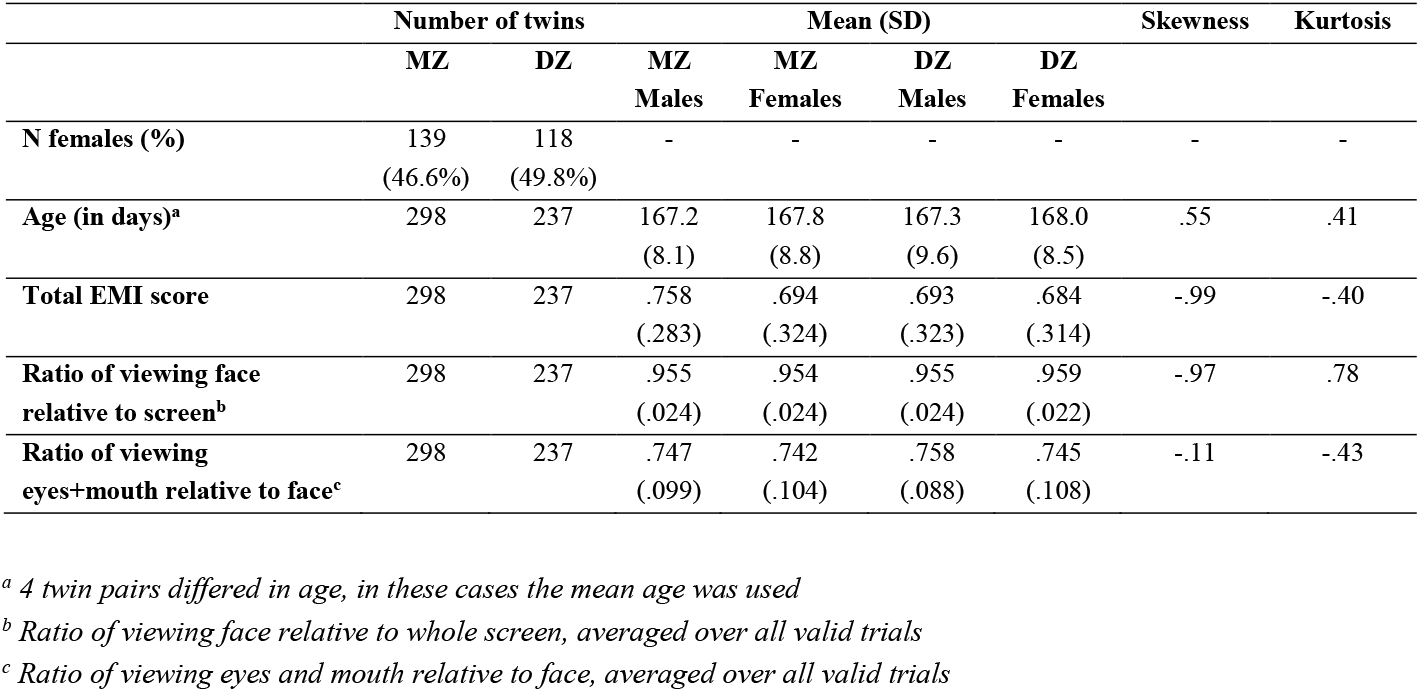
Descriptive statistics.

|  | Number of twins |  | Mean (SD) |  |  |  | Skewness | Kurtosis |
| --- | --- | --- | --- | --- | --- | --- | --- | --- |
|  | MZ | DZ | MZ | MZ | DZ | DZ |  |  |
|  |  |  | Males | Females | Males | Females |  |  |
| N females (%) | 139<br>(46.6%) | 118<br>(49.8%) | - | - | - | - | - | - |
| Age (in days)<sup>a</sup> | 298 | 237 | 167.2<br>(8.1) | 167.8<br>(8.8) | 167.3<br>(9.6) | 168.0<br>(8.5) | .55 | .41 |
| Total EMI score | 298 | 237 | .758<br>(.283) | .694<br>(.324) | .693<br>(.323) | .684<br>(.314) | -.99 | -.40 |
| Ratio of viewing face relative to screen<sup>b</sup> | 298 | 237 | .955<br>(.024) | .954<br>(.024) | .955<br>(.024) | .959<br>(.022) | -.97 | .78 |
| Ratio of viewing eyes+mouth relative to face<sup>c</sup> | 298 | 237 | .747<br>(.099) | .742<br>(.104) | .758<br>(.088) | .745<br>(.108) | -.11 | -.43 |

<sup>a</sup> 4 twin pairs differed in age, in these cases the mean age was used
<sup>b</sup> Ratio of viewing face relative to whole screen, averaged over all valid trials
<sup>c</sup> Ratio of viewing eyes and mouth relative to face, averaged over all valid trials

**Table 2.** Univariate twin model of the eye-mouth-index.

| Model | -2LL | #<br>parameters | df | AIC | Comparison<br>model | $\Delta \chi^2$ | $\Delta df$ | *p* | A | C | E |
| --- | --- | --- | --- | --- | --- | --- | --- | --- | --- | --- | --- |
| Fully sat. | 199.46 | 12 | 523 | -846.54 | - | - | - | - | - | - | - |
| ACE | 202.76 | 6 | 529 | -855.24 | Fully sat. | 3.29 | 6 | .77 | .47 | .09 | .44 |
| **AE** | **203.01** | **5** | **530** | **-856.99** | **ACE** | **0.25** | **1** | **.61** | **.57** | **-** | **.43** |
| CE | 209.21 | 5 | 530 | -850.79 | ACE | 6.45 | 1 | .01 | - | .45 | .55 |
| E | 263.27 | 4 | 531 | -798.73 | ACE | 60.51 | 2 | <.01 | - | - | 1 |

-2LL: Fit statistic, the lower the better fitting is the model
df: Degrees of freedom
AIC: An alternative fit index, lower value denotes better model fit
$\Delta \chi^2$ : Difference in $-2LL$ statistic between two models, distributed $\chi^2$
$\Delta df$ : Difference in degrees of freedom between two models

The eye-tracking procedure was conducted during the initial 5-month lab visit (Falck-Ytter et al., 2021). During the visit, the twins performed different tasks at the same time, in separate rooms. Several parent-report measures were administered again at 14 months as part of the larger aims of the study to track development.

### Measures

#### Eye tracking

Gaze data was recorded using the Tobii T120 Eye-tracker with a sampling rate of 60 Hz, using a standard Tobii monitor at native resolution (1024 x 768). The infant was seated in a baby chair or in the parent’s lap, approximately 60 cm from the screen. Before the eye tracking session, a 5-point calibration video was presented, and the experimental task did not begin until a successful calibration was achieved. Another 5-point video for offline calibration validation purposes was shown once in the beginning of the eye-tracking session.

For the main eye tracking analysis, each infant viewed 20 stimuli videos in a pseudo-random order. The videos comprised three conditions: Singing (twelve videos of a woman singing common Swedish nursery rhymes); Talking (four videos of a woman saying common Swedish rhyme verses); and Still (four videos of a woman smiling). In all videos, a woman was centered in the video and the background was grey (there were two women, each of them contributing equally to all conditions). The length of the videos ranged from 4 to 12 seconds.

Data were analyzed using custom scripts written in MATLAB (available upon request). Because the dependent measures were measures of accumulated looking time, we did not apply fixation filters (Kennedy et al., 2017). After data collection was finished, the data from the additional 5-point calibration video were evaluated via ocular inspection, and a simple linear transformation of data was performed when linear drifts were detected (using custom MATLAB scripts). Then, the total amount of looking time at the screen for each trial was calculated. Areas of interest (AOIs, i.e., face, eyes, and mouth) were created to move dynamically in coordination with the stimuli (using custom scripts in MATLAB), and were validated using visual inspection of their coordinates superimposed on the video stimuli. The face AOI was an ellipse with a horizontal radius of 200 pixels and a vertical radius of 280 pixels. Both the mouth AOI and eyes AOI were rectangles, 200×100 pixels and 310×100 pixels, respectively (https://www.smasyskon.se/stimuli).

Our primary dependent variable was the eye-mouth-index (EMI), which was calculated as the mean amount of gaze in the eyes AOI, divided by the mean amount of gaze to both the eyes AOI and the mouth AOI (i.e., 1 = only eyes looking; 0 = only mouth looking). The reason for using data from both the eyes and mouth region in the same metric (rather than separate them) is partly due to the use of the eye-mouth-index in earlier studies (Young et al., 2009). In addition, the eye-mouth-index is independent of differences in total gaze or stimuli duration. Constantino and colleagues (Constantino et al., 2017) did not report the EMI in their study, but instead reported separately preference for eyes and preference for mouth (each relative to the whole screen). Because infants tend to look predominantly at the eyes and the mouth when looking at a face, these two measures are highly and inversely correlated; hence, the current eye-mouth-index can be seen as largely analogous to the information reflected in the two measures reported in Constantino et al. (Constantino et al., 2017). Indeed, Constantino and colleagues reported that looking time in these two areas represented around 80% of infants’ looking time to the stimuli when observing videos with a single face included (the type of stimulus used for heritability estimation in their study). To verify that the infants focused mostly on the mouth and eyes region (instead of other regions of the face, e.g., chin or nose) we created an aggregated heatmap of gaze data inside the face AOI from all infants for all trials, which shows that the eyes and mouth were the primary regions of interest (https://www.smasyskon.se/stimuli). Further, we concluded that the eyes and mouth AOIs combined made up 75% of the gaze data towards the face (**Table 1**). See **Fig. S1** for the distributional properties of the EMI.

We then implemented steps to exclude trials based on general distribution properties. Specifically, using the results from infants with good calibration data (including after linear transformation), we obtained the values for the 10^th^ percentile for time spent looking at the screen, the 10^th^ percentile for the ratio of looking at the face (relative to the screen) and the 15^th^ percentile for the ratio of looking at the eyes and mouth combined (relative to the face) for each trial. If on a particular trial a participant was below one of these cut-offs or looked at the screen for a total of less than 1000 milliseconds, the trial was considered to be invalid, regardless of the classification of the calibration. Infants with at least four valid trials (from any condition) were included in further analyses. The number of included trials was not significantly associated with the EMI (r = .017, p = .699, N = 535).

In all three conditions, on group level, the infants preferred looking at the eyes (mean EMI singing condition = .71, SD = .32; mean EMI talking condition = .67, SD = .33; mean EMI still condition = .75, SD = .31). Because the phenotypic correlations between the three conditions were high (.871 – .925), we created one variable consisting of data from all infants with at least four valid trials (from any condition) and used this variable in all further analyses.

#### Parent-rated questionnaires

At 5 months, parents filled in the questionnaire version of the Vineland Adaptive Behavior Scales (Sparrow & Cicchetti, 1985). It is a standardized measure of adaptive behaviors across four domains. We used the standard scores for Communication and Socialization to measure socio-communicative behaviors at a concurrent age as the eye-mouth-index task.

The MacArthur Communicative Development Inventory, CDI (Fenson et al., 1993). is a parent-rated questionnaire that assesses early language development and was administered at 14 months (the Words and Gestures form) and 24 months (the Words and Sentences form). As a measure of expressive vocabulary at 14 months, we used the total raw word production score, which is the number of words (out of 370 words) that the infant can produce. Parents also reported whether the infant could understand but not produce the words, which is summarized as the total comprehension score. This score was used as a measure of receptive vocabulary. At 24 months, we used the vocabulary checklist score as a measure of expressive vocabulary.

The CSBS DP Infant Toddler Checklist, ITC (Wetherby & Prizant, 2002), is a 24-item parent-rated questionnaire, used to identify children with any type of communication delay, including ASD. Lower scores indicate a higher degree of socio-communicative delays. It was administered at 14 months, and we used the total score as a measure of socio-communicative behaviors linked to ASD.

The Quantitative Checklist for Autism in Toddlers, Q-CHAT (Allison et al., 2008), is a normally distributed quantitative measure of autistic traits, which consists of 25 parent-rated items scored on a 5-point scale (0-4) and was administered at 24 months. The scores from all items are summed to obtain a total score, where higher scores indicate more autistic traits.

#### Experimenter-rated developmental assessment

The Mullen Scales of Early Learning, MSEL (Mullen, 1995), was administered by an experimenter at 5 months. This is a standardized assessment commonly used in many areas of psychology as a measure of cognitive development. The MSEL consists of five subscales (gross motor, fine motor, visual reception, receptive language, and expressive language). See **Table S2** for descriptive statistics on parent-rated questionnaires and the experimenter-rated developmental assessment.

### Statistical analyses

An analysis plan was pre-registered in OSF (https://osf.io/s8y74/) after data collection but prior to data analysis. We used a univariate twin model to estimate the genetic and environmental contribution to variation in eye-mouth-index. The sources of variation in a trait can be divided into genetic influences (A; heritability), shared environment (C; e.g., family environment), and unique environment (E; i.e., environmental influences that makes twins different from each other, including measurement error). Since monozygotic (MZ) twins share 100% of their segregating DNA, while dizygotic (DZ) twins on average share 50% of their segregating DNA, a higher within pair similarity among MZ twin than DZ twins suggests genetic contribution to a trait. The overall EMI score was used for the univariate twin model as well as all further analyses. Sex and age were incorporated as covariates. Data analysis was performed in R 3.6.3 (R Team, 2017), and model fitting was performed through maximum likelihood optimization with OpenMx, version 2.17.2 (Neale et al., 2016).

Associations between EMI at 5 months and vocabulary at 14 months using the CDI and socio-communicative behaviors at 14 months using the ITC were calculated using the robust sandwich estimator in generalized estimating equations (GEE) in order to account for the correlation between twins in a pair (Carlin, Gurrin, Sterne, Morley, & Dwyer, 2005). The variables used in these phenotypic associations were regressed on age and sex before further analyses.

## Results

### Genetic analyses

The twin correlations for the EMI score were higher for monozygotic twins than dizygotic twins (rMZ = .55, 95% CI: .42, .65; rDZ = .34, 95% CI: .16, .49), suggesting genetic influence. A fully saturated model was fitted in order to test the assumptions of equality of means and variances across zygosity and twin order (see **Table S3** for details). Based on the twin correlations, we fitted an ACE model, along with AE, CE, and E models for comparison. Based on the likelihood-ratio test and the AIC value, the best fitting model was an AE model, where the shared environment component was dropped. The AE model’s estimates suggested a moderate-to-high heritability of the preference for looking at eyes versus mouth (A = .57; 95% CI: .45, .66), with a moderate contribution of non-shared environment (E = .43; 95% CI: .34, .55). Further, we tested the association between EMI and polygenic scores for autism spectrum disorder, educational attainment, and IQ. None of these associations were statistically significant (**Table S1**).

### Longitudinal phenotypic associations

Contrary to the hypothesis, a trend towards a *positive* association was found between the EMI and the CDI production score (β = .10; 95% CI: −.01, .21; p = .064; N = 419). However, the production score from the CDI had a considerable floor effect, reflecting the fact that most infants produced none or only a few words. Therefore, in a deviation from the pre-registered plan, we analyzed the CDI comprehension score as well, which showed a statistically significant positive association with EMI (β = .16; 95% CI: .05, .27; p < .01; R^2^ = .03; N = 419). No significant association was found between autistic traits (the total score on ITC) and the EMI (β = .08; 95% CI: −.03, .19; p = .175; N = 418).

### Secondary phenotypic analyses

In light of the results above, we conducted a series of follow up analyses, to probe the degree of independence of the EMI measure from other concurrent developmental domains, and of the specificity of the association between EMI and language development at 14 months. First, we tested the association with expressive vocabulary at 24 months (this time point was excluded from the original pre-registration due to the increasing attrition rate with age). This measure was not associated with the EMI at 5 months (β = .02; 95% CI: −.11, .15; p = .783; N = 341; CDI at 24 months does not include a comprehension score), suggesting that the link may be specific to parental ratings of verbal competence in early toddlerhood. Second, we followed up the negative finding regarding autistic traits at 14 months (ITC) by analyzing scores on the Quantitative Checklist for Autism in Toddlers (Q-CHAT) at 24 months, and again, no significant association was detected (β = −.02; p = .696; N = 343). Next, we tested whether the EMI was associated with general development and socio-communicative behavior at 5 months using the Mullen Scales of Early Learning (MSEL; a standardized assessment of cognitive development consisting of five subscales: gross motor, fine motor, visual reception, receptive language, and expressive language) and the Vineland Adaptive Behavior Scales (a standardized measure of adaptive behavior from which we used the Communication domain and the Socialization domain). We found no significant association between the EMI and any of these concurrent measures, suggesting that EMI is a highly independent heritable phenotype in early infancy (**Table S4**).

Finally, we tested the specificity of the association between EMI and 14-month language comprehension in light of the other concurrent 5-month measurements available. A GEE analysis with multiple predictors showed that experimenter ascertained gross motor ability (MSEL gross motor scale at 5 months), parental ratings of social skills (Vineland Socialization scale at 5 months) and the EMI (5 months) all had unique contributions to the CDI comprehension ratings at 14 months (**Table 3**).

**Table 3.** A GEE analysis with receptive vocabulary (language comprehension) at 14 months as outcome variable, including eight predictors measured at 5 months (n = 401).

|  | **Beta** | **95% CI** | **p** |
| --- | --- | --- | --- |
| **EMI** | .13 | .03; .22 | **.011** |
| **MSEL** |  |  |  |
| Gross motor | .12 | .01; .23 | **.040** |
| Visual reception | .08 | -.03; .18 | .138 |
| Fine motor | .02 | -.08; .12 | .672 |
| Receptive language | .01 | -.08; .10 | .816 |
| Expressive language | .01 | -.09; .11 | .897 |
| **Vineland** |  |  |  |
| Communication | .12 | -.00; .24 | .059 |
| Socialization | .19 | .06; .32 | **.003** |

*CDI: Communicative Development Inventory (Words and Gestures form)*
*EMI: Eye-Mouth-Index*
*MSEL: Mullen Scales of Early Learning*
*Vineland: Vineland Adaptive Behavior Scales*

## Discussion

This study demonstrates that individual differences in young infants’ preference for specific parts of others’ faces (specifically the preference for eyes versus mouth, EMI) to a large extent can be explained by genotypic differences. The contribution of non-shared environment (which also includes error of measurement) was moderate, while shared environment does not appear to influence infant social viewing in this context. Our finding indicates that before infants can select their environment by means of crawling or walking, they select aspects of the environment that they look at largely based on their individual genotypes; i.e. a type of active gene-environment correlation operating at short time scales via gaze behavior (Kennedy et al., 2017).

We found that eye versus mouth preference was highly correlated (~.9) across conditions. Thus, it appears that individual differences in EMI at this age were largely independent of the exact stimulus properties such as movement and audio-visual synchrony (that differed between the conditions), a finding which is in line with the patterns of results observed by Constantino et al. in toddlers (Constantino et al., 2017). Possibly, infants may use basic invariant face properties as a basis for their preference, such as face-like configurations “eyes over mouth”, to which infants are sensitive from very early on (Quinn & Tanaka, 2009; Schwarzer, Zauner, & Jovanovic, 2007).

Eye preference did not correlate with other concurrent traits assessed in the study, including developmental domains directly assessed by the experimenter (motor, perceptual, communication) as well as parent-rated skills (social communication). This suggests that eye versus mouth preference at five months reflect a highly independent, and strongly heritable, socio-attentional trait.

We found that preference for the eyes at five months was positively correlated with parents’ assessment of vocabulary at 14 months, measured with the CDI. This result is somewhat surprising given previous research showing an association between large vocabulary and a preference for looking at the mouth (Tenenbaum et al., 2015; Young et al., 2009). It is notable, however, that we assessed EMI at slightly earlier age compared to previous reports. At around five months, infants learn to follow other people’s gaze (Del Bianco, Falck-Ytter, Thorup, & Gredeback, 2019), which is known to facilitate word learning (by following gaze, infants will attend to the same things as their caregivers, and hence more quickly understand what they refer to when they name objects and events). Thus, our results indicate that at this early age, attending to cues relevant for joint attention is more important for later language, than attending to speech cues linked to the mouth.

It is notable that eye preference predicted parents’ ratings of language comprehension over and above variance captured by other scales at 5 months, suggesting high specificity. Further, eye looking seems to specifically predict ratings of language ability in early toddlerhood, while associations with ratings at 24 months were non-significant. This pattern is consistent with the concept of equifinality in development (Cicchetti & Rogosch, 1996): while infants who attend to the mouth rather than eyes may have a temporary disadvantage in terms of vocabulary development, over time they catch up. Such diversity in developmental pathways (which may be differentially adaptive in different contexts) may also help explain why the genetic variance has remained in the population.

A limitation of our protocol is the reliance on parental ratings of language development in toddlerhood and early childhood. However, the CDI scores have been shown to be stable over time and associated with later language ability (Berglund & Eriksson, 2000; Eriksson, 2001; Fenson et al., 1993, but see Houston-Price, Mather, & Sakkalou, 2007). In addition to differences in true ability, CDI ratings may reflect parents’ *perception* of ability. Infants’ eye movements are visible to others – where they look in the face may also influence their parents’ impression of them (Constantino et al., 2017; Kennedy et al., 2017). Nevertheless, the results are important because they demonstrate that a heritable trait in infancy predict how caregivers perceive their offspring’s competence almost a year later.

The lack of association between the EMI and later socio-communication difficulties is consistent with the results of Constantino et al. (2017) who found that toddlers with ASD did not have an atypical preference for the eyes versus the mouth, but rather looked less at both these areas when observing complex scenes with multiple competing objects (the ASD sample was only tested using complex stimuli with multiple people and objects). What may be most pronounced in ASD (or in individuals with high autistic traits) is not atypical eye *versus* mouth preference, but reduced attention to faces overall (Dewaele, Demurie, Warreyn, & Roeyers, 2015; Guillon et al., 2016; Shic, Macari, & Chawarska, 2014). To the best of our knowledge, no study has yet investigated heritability of face preference at any age. Thus, this should be a high priority for future studies.

The high heritability and specificity of the eye (vs mouth) preference found at five months in this study raises the question of to what extent other brain and behavioural traits are equally heritable early in life, and to what extent those genetic factors are general or phenotype specific. Our results point to the promise of combining genetically informed designs with state-of-the-art infant research technology.

## Acknowledgments

The authors thank all participating families, as well as researcher Pär Nyström and research assistants Linnea Hamrefors, Joy Hättestrand, Lynnea, Myers, Johanna Kronqvist, Sofia Jönsson, Anna Kernell, Carolin Schreiner, Sophie Lingö, Angelinn Liljebäck, Isabelle Enedahl, Matthis Andreasson, Lisa Belfrage, Mattias Savallampi, Isabelle Ocklind and Hjalmar Nobel Norrman. The genotyping was done at the SNP&SEQ Technology Platform, Uppsala University.

## Funding

Swedish Research Council, grant 2018-06232 (TFY)

Riksbankens Jubileumsfond in collaboration with the Swedish Collegium for Advanced Study (TFY)

Knut and Alice Wallenberg Foundation (TFY)

Innovative Medicines Initiative 2 Joint Undertaking under grant agreement No 777394. This Joint Undertaking receives support from the European Union’s Horizon 2020 research and innovation programme and EFPIA and AUTISM SPEAKS, Autistica, SFARI. (TFY)

## Author contributions

The hypotheses and goals of this study were conceptualized by C.V, A.M.P, M.R., P.N, A.R. and T.F-Y. Data were analyzed by C.V, with input from A.M.P, M.J.T, A.R, M.S.S, and T.F-Y. Software was programmed by P.N, M.J.T, and C.V. Polygenic scores were derived by D.L and K.T. The research was supervised by T.F-Y. C.V and T.F-Y drafted the manuscript, and all of the authors reviewed, edited, and approved the final manuscript for submission.

## Data and materials availability

Available upon reasonable request to corresponding author. Note that sharing of pseudonymized personal data will require a data processor agreement (DPA), according to Swedish and EU law.

