## Supplementary Information for "Infants’ attention to eyes is an independent, heritable trait predicting later verbal competence"

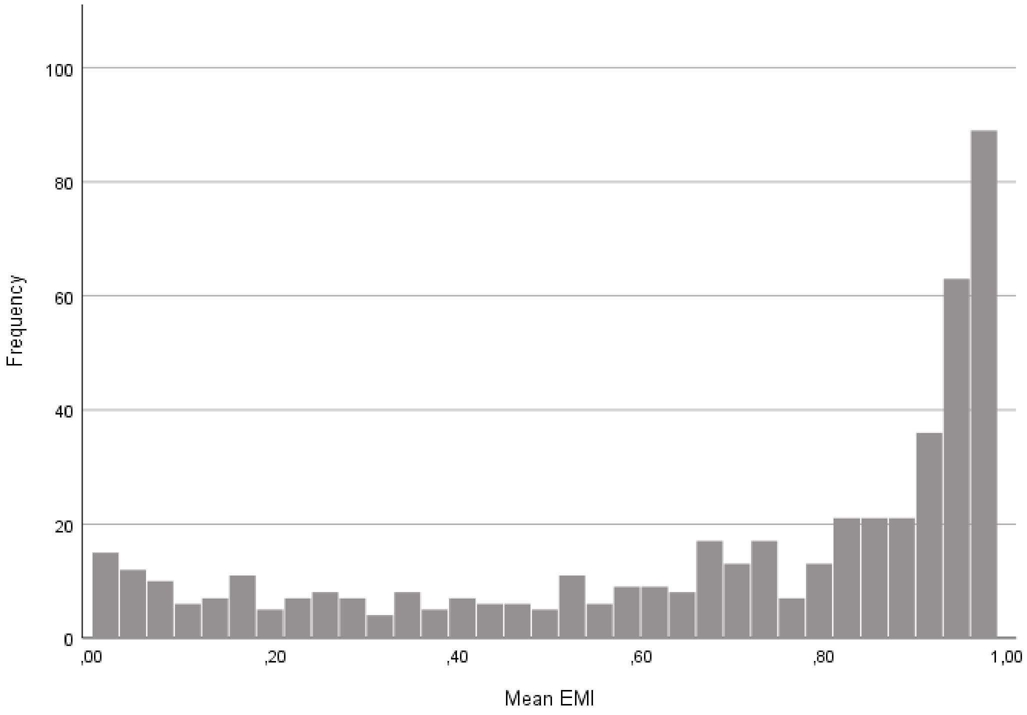


**Fig. S1.**

Distribution of the EMI variable (N = 535 infants).

GEE analyses with EMI as the dependent variable and polygenic scores as predictors

| | Beta | 95% CI | p | n |
|---|---|---|---|---|
| **Polygenic score ASD** | .00 | -.04; .04 | .986 | 527 |
| **Polygenic score EA** | -.01 | -.05; .03 | .580 | 527 |
| **Polygenic score IQ** | -.02 | -.06; .01 | .152 | 527 |

**Table S1. Phenotypic analyses between EMI and polygenic scores for autism spectrum disorder (ASD), educational attainment (EA), and IQ**. Due to the association between mouth-looking and vocabulary, which in turn predicts school achievement (Bleses, Makransky, Dale, Hojen, & Ari, 2016), we tested whether more looking at the mouth was associated with higher polygenic scores related to cognition (i.e., IQ and educational attainment). We also tested the association between eye versus mouth looking and polygenic scores for ASD, but as for the phenotypic association, no directional hypothesis was formulated.

The use of polygenic scores requires a discovery dataset and a target dataset. The discovery dataset is used to identify individual SNPs (single nucleotide polymorphisms) that are associated with the trait at a statistically significant level. Results from the discovery dataset are then used to derive polygenic scores in the target dataset by creating a weighted sum of all the alleles that either increase or decrease the chance of the outcome of interest. As discovery datasets for this study, we used the results of the genome-wide association study (GWAS) of IQ carried out by the Complex Trait Genetics lab (Savage et al., 2018), the GWAS of educational attainment by the Psychiatric Genomics Consortium (Lee et al., 2018), and the GWAS of ASD by the Psychiatric Genomics Consortium (Grove et al., 2019). Our current sample of 5 months old twins was used as the target dataset. In summary, the polygenic scores were calculated using the PRS-CS method (Ge, Chen, Ni, Feng, & Smoller, 2019), but details are reported elsewhere (Falck-Ytter et al., 2021). Three separate GEE analyses were performed to assess the association between polygenic scores and EMI. Sex, age, and the first 10 principal components of ancestry were included as covariates.

*EMI: Eye-Mouth-Index*

Phenotypic analyses: descriptive statistics

| | Number of infants | Mean (SD) | Skewness | Kurtosis |
|---|---|---|---|---|
| **MSEL 5 months** | | | | |
| Gross motor | 559 | 7.88 (1.14) | -.14 | .92 |
| Visual reception | 560 | 7.18 (.97) | .19 | 7.90 |
| Fine motor | 557 | 6.73 (1.22) | .29 | -.07 |
| Receptive language | 558 | 6.33 (1.05) | -.22 | 1.21 |
| Expressive language | 558 | 6.06 (.77) | -.13 | 1.19 |
| **Vineland* 5 months** | | | | |
| Communication | 567 | 39.96 (2.66) | -2.00 | 7.92 |
| Socialization | 567 | 50.70 (2.44) | 1.13 | .28 |
| **CDI 14 months (production)** | | | | |
| Females | 201 | 8.17 (21.64) | 8.75 | 88.30 |
| Males | 218 | 6.11 (10.60) | 5.12 | 36.32 |
| **CDI 14 months (comprehension)** | | | | |
| Females | 201 | 94.53 (68.50) | 1.34 | 2.70 |
| Males | 218 | 74.74 (66.52) | 1.47 | 2.01 |
| **ITC 14 months** | | | | |
| Females | 200 | 35.88 (6.14) | -.19 | .15 |
| Males | 218 | 34.00 (7.76) | -.34 | -.21 |
| **CDI 24 months** | | | | |
| Females | 186 | 250.19 (144.53) | .67 | -.23 |
| Males | 175 | 168.07 (163.56) | 1.15 | .48 |
| **Q-CHAT 24 months** | | | | |
| Females | 191 | 24.66 (6.80) | -.46 | .73 |
| Males | 172 | 29.26 (8.01) | .51 | .44 |

*The standard scores have been derived from norms for 2-year-olds, due to it being the lowest age for which there exists Swedish norms

### Table S2. Descriptive statistics for the variables included in the phenotypic analyses.

*MSEL: Mullen Scales of Early Learning*

*Vineland: Vineland Adaptive Behavior Scales*

*CDI: Communicative Development Inventory*

*ITC = Infant Toddler Checklist*

*Q-CHAT: The Quantitative Checklist for Autism in Toddlers*

Testing covariates and assumptions

| | | | | | | Comparative fit with saturated model | | |
|---|---|---|---|---|---|---|---|---|
| **Model** | **-2LL** | **# parameters** | ***df*** | **AIC** | **BIC** | **Δ χ2** | **Δ df** | **p** |
| Fully sat. | 199.46 | 12 | 523 | -846.54 | -2764.08 | - | - | - |
| Submodel 1 | 200.37 | 10 | 525 | -849.63 | -2774.50 | 0.91 | 2 | .63 |
| Submodel 2 | 200.71 | 8 | 527 | -853.29 | -2785.50 | 1.25 | 4 | .87 |
| Submodel 3 | 202.22 | 7 | 528 | -853.78 | -2789.65 | 2.76 | 5 | .74 |
| Submodel 4 | 202.76 | 6 | 529 | -855.24 | -2794.78 | 3.29 | 6 | .77 |
| Age | 200.77 | 11 | 524 | -847.23 | -2768.44 | 1.30 | 1 | .25 |
| Sex | 200.54 | 11 | 524 | -847.46 | -2768.67 | 1.08 | 1 | .30 |

**Table S3**. **Analyses of covariates and assumptions for the univariate twin model of EMI.** The fully saturated model is the baseline model, which models the means and variances separately for each twin in a pair and across zygosity.

*Submodel 1: Equating means across twins within a pair*
*Submodel 2: Equating means across zygosity*
*Submodel 3: Equating variances across twins within a pair*
*Submodel 4: Equating variances across zygosity*
*Age: Testing the significance of age*
*Sex: Testing the significance of sex*
*-2LL: Fit statistic, the lower the better fitting is the model*
*df: Degrees of freedom*
*AIC: An alternative fit index, lower value denotes better model fit*
*BIC: An alternative fit index, lower value denotes better model fit*
*Δ χ2: Difference in −2LL statistic between two models, distributed χ2*
*Δ df: Difference in degrees of freedom between two models*

GEE analyses with EMI as the dependent variable

| | **Beta** | **95% CI** | **p** | **n** |
|---|---|---|---|---|
| **MSEL** | | | | |
| Gross motor | .05 | -.05; .14 | .352 | 521 |
| Visual reception | -.06 | -.16; .04 | .242 | 522 |
| Fine motor | -.03 | -.13; .06 | .483 | 519 |
| Receptive language | -.01 | -.11; .09 | .812 | 520 |
| Expressive language | -.01 | -.10; .08 | .787 | 520 |
| **Vineland** | | | | |
| Communication | -.01 | -.09; .08 | .846 | 528 |
| Socialization | .01 | -.10; .13 | .819 | 528 |

**Table S4. Phenotypic analyses of EMI and MSEL/Vineland**. All phenotypic analyses were calculated using the robust sandwich estimator in generalized estimating equations (GEE) in order to account for the correlation between twins in a pair. The variables were regressed on age and sex before analysis.

*EMI: Eye-Mouth-Index*
*MSEL: Mullen Scales of Early Learning*
*Vineland: Vineland Adaptive Behavior Scales*
